# Iron-loaded deferiprone can support full hemoglobinization of cultured red blood cells in the absence of transferrin

**DOI:** 10.1101/2021.08.02.454758

**Authors:** Joan Sebastián Gallego-Murillo, Nurcan Yağcı, Eduardo Machado Pinho, Adrián Abeijón-Valle, Aljoscha Wahl, Emile van den Akker, Marieke von Lindern

**Author notes:** Correspondence: Marieke von Lindern, Department of Hematopoiesis, Sanquin Research and Landsteiner Laboratory, Amsterdam UMC, Plesmanlaan 125, 1066CX Amsterdam, The Netherlands; and Joan Sebastián Gallego-Murillo, Department of Hematopoiesis, Sanquin Research and Landsteiner Laboratory, Amsterdam UMC, Plesmanlaan 125, 1066CX Amsterdam, The Netherlands;.

## Abstract

Iron is an essential nutrient in mammalian cell cultures, conventionally supplemented as iron-loaded transferrin (holotransferrin). The high cost of human transferrin represents a challenge for the large scale production of cell therapies, such as cultured red blood cells. We evaluated the use of deferiprone, a cell membrane-permeable drug for iron chelation therapy, as an iron carrier for erythroid cultures. Iron-loaded deferiprone (Def_3_·Fe^3+^) at a concentration of 52μmol/L could fully replace holotransferrin during erythroblast differentiation into reticulocytes, the erythroid differentiation stage with maximal iron requirements. Reticulocytes cultured in presence of Def_3_·Fe^3+^ or holotransferrin (1000μg/mL) were similar with respect to expression of cell-surface markers CD235a and CD49d, hemoglobin content, and oxygen association/dissociation. Def_3_·Fe^3+^ also supported expansion of the erythroid compartment *in vitro*, except for the first stage when hematopoietic stem cells committed to erythroblasts, in which a reduced erythroblasts yield was observed. This suggests that erythroblasts acquired the potential to process Def_3_·Fe^3+^ as iron source for biosynthesis pathways. Replacement of holotransferrin by Def_3_·Fe^3+^ was also successful in cultures of six myeloid cell lines (MOLM13, NB4, EOL1, K562, HL60, ML2). These results suggest that iron-loaded deferiprone can partially replace holotransferrin in chemically defined medium formulations for the production of cultured reticulocytes and proliferation of selected myeloid cell lines. This would lead to a significant decrease in medium cost that would improve the economic perspectives of the large scale production of red blood cells for transfusion purposes.

**Key points:**

- Holotransferrin limitations in erythroid cultures lead to lower erythroblast yields, impaired maturation and low enucleation efficiencies.
- Iron-loaded deferiprone can replace holotransferrin in erythroblast expansion and differentiation cultures.

## Introduction

Iron is an essential nutrient for all eukaryotic microorganisms. Erythroid precursors, however, have exceptionally high iron requirements to facilitate hemoglobin synthesis during the generation of red blood cells (RBCs). A single RBC contains approximately 300 million molecules of hemoglobin (≈30pg), associated with 1.2×10^9^ Fe^3+^ ions.^1^ In healthy humans, 65-75% of all body iron is present as hemoglobin in RBCs.^2^ Free ferrous iron Fe^2+^ can lead to the production of toxic radicals via the Fenton reaction.^3^ Therefore, plasma iron is transported bound to e.g. transferrin (Tf). Human transferrin is a single chain glycoprotein with a molecular weight of ~80kDa. It has two distinct iron-binding lobes (N- and C-lobe). The binding of Fe^3+^ to Tf is sequential, resulting in four possible Tf species: iron-free Tf (aTf: apotransferrin), diferric Tf (hTf: holotransferrin), and two monoferric Tf forms depending on which lobe contains the Fe^3+^ ion.^4^

High expression of the transferrin receptor (TfR; CD71) on erythroblasts facilitates their increased uptake of hTf for heme synthesis. Iron deficiency reduces the yield of burst-forming erythroid-units in early stages of *ex vivo* erythropoiesis,^5^ and of reticulocytes during terminal erythroid differentiation.^6^ Assuming that hTf is the sole source of iron, ~80pg of hTf needs to be internalized for a single fully-hemoglobinized RBC, corresponding to ~160g hTf for a single transfusion unit of RBCs (2×10^12^ RBCs). Transferrin is endocytosed upon binding to TfR. Fe^3+^ ions are released at the low endocytic pH, reduced to Fe^2+^ by STEAP3, transported into the cytosol and subsequently to mitochondria to be incorporated into heme.^7,8^ Excess iron is bound to cytosolic ferritin.^9^ Vesicles containing aTf still bound to TfR recycle to the cell surface where aTf is released. In the body, the released aTf can bind new Fe^3+^ ions via direct transfer from ceruloplasmin ferroxidase.^10^

The expression level of key regulatory proteins involved in iron uptake, storage and export is controlled by the RNA-binding proteins Iron regulatory protein-1 and −2 (IRP1/2). IRP1/2 binding to Fe-S clusters changes its conformation, and inhibits binding to iron-responsive elements (IREs) in the untranslated regions of mRNAs. For iron uptake proteins such as TfR and DMT1, IRP1 binding to IREs in the 3’ UTR of respective mRNAs increases mRNA stability. By contrast, IRP1 binding to the IREs in the 5’ UTR of mRNAs encoding proteins involved in iron storage and export (e.g. ferritin and ferroportin) decreases their translation.^11,12^ Iron deficiency decreases heme levels, which activates Heme-regulated eIF2α kinase (HRI), resulting in phosphorylation of eIF2 and a global inhibition of protein translation.^13(p2),14^ At the level of erythropoiesis, lack of iron causes microcytic anemia in vivo,^15^ and decreases the proliferation and differentiation of erythroblast cultures *ex vivo*.^6^

In serum-free cell culture systems, high levels of hTf (1000μg/mL) are added during terminal differentiation.^16,17^ Currently, most of the Tf used in cultures is purified from human plasma. However, there is a growing need to transition towards animal-free media components to ensure safety of the final product. Production of recombinant Tf in *Escherichia coli*, insects, yeast, rice and BHK cells represents an alternative, but still not cost-effective for the *ex vivo* production of RBCs.^18^

Several small molecules have been reported to chelate and deliver iron into the cells. PIH (pyridoxal isonicotinoyl hydrazine) restored proliferation and differentiation in several cell lines in absence of hTf by directly crossing the cellular membrane.^19,20^ Supplementation of hinokitiol was also shown to restore proliferation and hemoglobinization in DMT1-deficient cells in transferrin-free medium, although its effectiveness seems to depend on large iron gradients across the cellular membrane.^21^

Compounds with a high affinity for iron that are commonly used to treat iron overload *in vivo* have also been tested in cell culture. Among them is deferiprone (1,2-dimethyl-3-hydroxypyridin-4-one), a bidentate alpha-keto hydroxypyridine molecule that binds iron with a stoichiometry of 3:1 (Def_3_·Fe^3+^) at physiological conditions.^22^ Its ability to cross the cell membrane and mobilize intracellular iron is the principle of combined chelation therapy, in which deferiprone scavenges intracellular iron and transfers it to a second non-permeable chelator such as deferoxamine.^23,24^

In this study, we evaluated the potential of iron-loaded deferiprone (Def_3_·Fe^3+^) to replace hTf in cultures of erythroid precursors for the *ex vivo* production of RBCs. We demonstrated that Def_3_·Fe^3+^ can recharge aTf to hTf. However, medium supplementation with Def_3_·Fe^3+^ alone was sufficient to sustain efficient *ex vivo* production of fully hemoglobinized RBCs with an oxygen binding capacity comparable to peripheral blood RBCs. Excess Def_3_·Fe^3+^ showed no sign of toxicity in erythroblast proliferation and differentiation. Additionally, the use of Def_3_·Fe^3+^ as the only iron source in serum-free medium was sufficient to sustain the proliferation of other transferrin-dependent mammalian cell lines.

## Materials and methods

### Cell culture

Donor-derived buffy coats are a Sanquin not-for-transfusion product from which PBMCs are purified by density centrifugation. Informed consent was given in accordance with the Sanquin Ethical Advisory Board. Erythroid cells were cultured from human PBMCs as previously described^25^ with minor modifications to Cellquin medium, which lacked nucleosides, and contained a defined lipid mix (Sigma-Aldrich; USA; 1:1000) replacing cholesterol, oleic acid, and L-α-phosphatidylcholine. Transferrin concentrations varied as indicated. Expansion cultures were supplemented with EPO (2U/mL; EPREX^®^, Janssen-Cilag, Netherlands), hSCF (100ng/mL, produced in HEK293T cells), dexamethasone (1μmol/L; Sigma-Aldrich), and IL-3 (1ng/mL first day only; Stemcell Technologies; Canada). Cell density was maintained between 0.7-2×10^6^ cells/mL (CASY Model TCC; OLS OMNI Life Science; Germany). To induce differentiation, cells were washed and reseeded at 1-2×10^6^ cells/mL in presence of EPO (10U/mL), 5% Omniplasma (Octapharma GmbH; Germany), heparin (5U/mL; LEO Pharma A/S; Denmark), and hTf (Sanquin; Netherlands) or aTf (Sigma-Aldrich) as indicated. AML-derived cell lines MOLM13,^26^ NB4,^27^ EOL1,^28^ K562,^29^ HL60,^30^ and ML2 ^31^ were seeded at a concentration of 0.3×10^6^ cells/mL in Cellquin, supplemented with hTf and Def_3_·Fe^3+^ as indicated.

### Iron chelators

Deferiprone (Def; Sigma-Aldrich), deferoxamine mesylate (DFOA; Sigma-Aldrich), deferasirox (DFO; Sigma-Aldrich) and hinokitiol (HINO; Sigma-Aldrich) were dissolved as indicated by the manufacturer and mixed at stoichiometric ratio with an iron(III) chloride solution (Sigma-Aldrich; 16h, 20°C), to obtain a concentration of 26mmol/L chelated Fe^3+^ (Def_3_·Fe^3+^, DFOA·Fe^3+^, DFO_2_·Fe^3+^, HINO_3_·Fe^3+^), equivalent to the iron content of 1000mg/mL hTf.

### Flow cytometry

Cells were stained in HEPES buffer + 0.5% BSA (25-30 minutes, 4°C), measured using a BD FACSCanto^™^ II flow cytometer (BD Biosciences), gated against specific isotypes, and analyzed using FlowJo™ (version 10.3; USA). Antibodies or reagents used were: (i) CD235a-PE (1:2500 dilution; OriGene cat#DM066R), CD49d-BV421 (1:100 dilution; BD-Biosciences cat#565277), DRAQ7 (live/dead stain; 1:200 dilution; ThermoFischer Scientific cat#D15106); (ii) CD235a-PE (1:2500 dilution; OriGene cat#DM066R), CD71-APC (1:200 dilution; Miltenyi cat#130-099-219); (iii) CD71-VioBlue (1:200 dilution; Miltenyi cat#130-101-631); (iv) DRAQ5 (nuclear stain; 1:2500 dilution; abcam cat#ab108410); (v) PI (live/dead stain; 1:2000 dilution; Invitrogen cat#P3566).

### Western blots

Iron saturation of transferrin was measured on 6% TBE-urea gels as previously described.^32^ Gels were stained with InstantBlue Coomassie protein stain (abcam).

Whole cell lysates were prepared in RIPA buffer (10min, 4°C). Protein concentration was determined by colorimetry (DC^™^ Protein Assay; Bio-Rad; USA). Lysates were diluted 1:4 with Laemmli sample buffer (Bio-Rad), incubated for 95°C (5min), subjected to SDS-polyacrylamide gel electrophoresis (4–20% Criterion^™^ Tris-HCl Protein Gel, Bio-Rad), transferred to nitrocellulose membranes (iBlot2 system; ThermoFischer Scientific), and stained. Primary antibodies included actin (Sigma cat#A3853), transferrin receptor (Merck cat#SAB4200398), ferritin (abcam cat#ab75973), ferroportin (abcam cat#ab235166), ALAS2 (Merck cat#SAB2100096), p-eIF2 (CellSignaling cat#3597L) and total eIF2 (CellSignaling cat#9722S). Western blots were analyzed using GelAnalyzer 19.1.

### Hemoglobin quantification and oxygenation

Hemoglobin content was determined using *o*-phenylenediamine as described.^33^ Intracellular hemoglobin concentration was calculated using the mean of technical triplicates and the cell volume (CASY Model TCC). Oxygen association/dissociation curves for RBCs were determined by continuous dual wavelength spectrophotometric measurement with a HEMOX Analyzer (TCS Scientific Corp.; USA), using RBCs that were washed and resuspended at 20×10^6^ cells/mL in analysis buffer (PBS + 0.5% BSA + 0.005% Y-30 antifoam).

### Data sharing statement

For original data, please contact.

## Results

### Iron-loaded deferiprone supports erythroid differentiation at low hTf concentrations

#### Reference conditions

As reference for the different iron supplementation strategies, we first tested different holotransferrin concentrations during erythroblast differentiation, the culture stage in which cells have the highest iron requirements. Twelve days post-seeding PBMCs in expansion medium, erythroblasts were cultured in medium supplemented with decreasing concentrations of hTf (1000, 200, 100, 50 and 0μg/mL; equivalent to 26, 5.2, 1.3 and 0μmol/L of chelated Fe^3+^ respectively). At 1000μg/mL hTf, erythroblasts proliferated for 3-4 days followed by cell growth arrest at the final stage of differentiation.^25^ A gradual decrease of hTf concentrations resulted in a gradual decrease in cell yields, in accordance with previously described data (Figure 1A).^6^ Erythroblast differentiation is accompanied by a progressive loss of cell volume to reach the cell size of erythrocytes. Suboptimal hTf concentrations led to a faster decrease in cell size compared with 1000μg/mL hTf until days 4-5, and a consistently lower cell volume that was still present after 8 days of culture (Supplemental Figure S1A). This is in accordance with microcytic anemia upon iron deficiency. Hemoglobin levels were low at the start of differentiation, and rapidly increased dependent on the hTf concentration (Supplemental Figure S1B).

**Figure 1.**
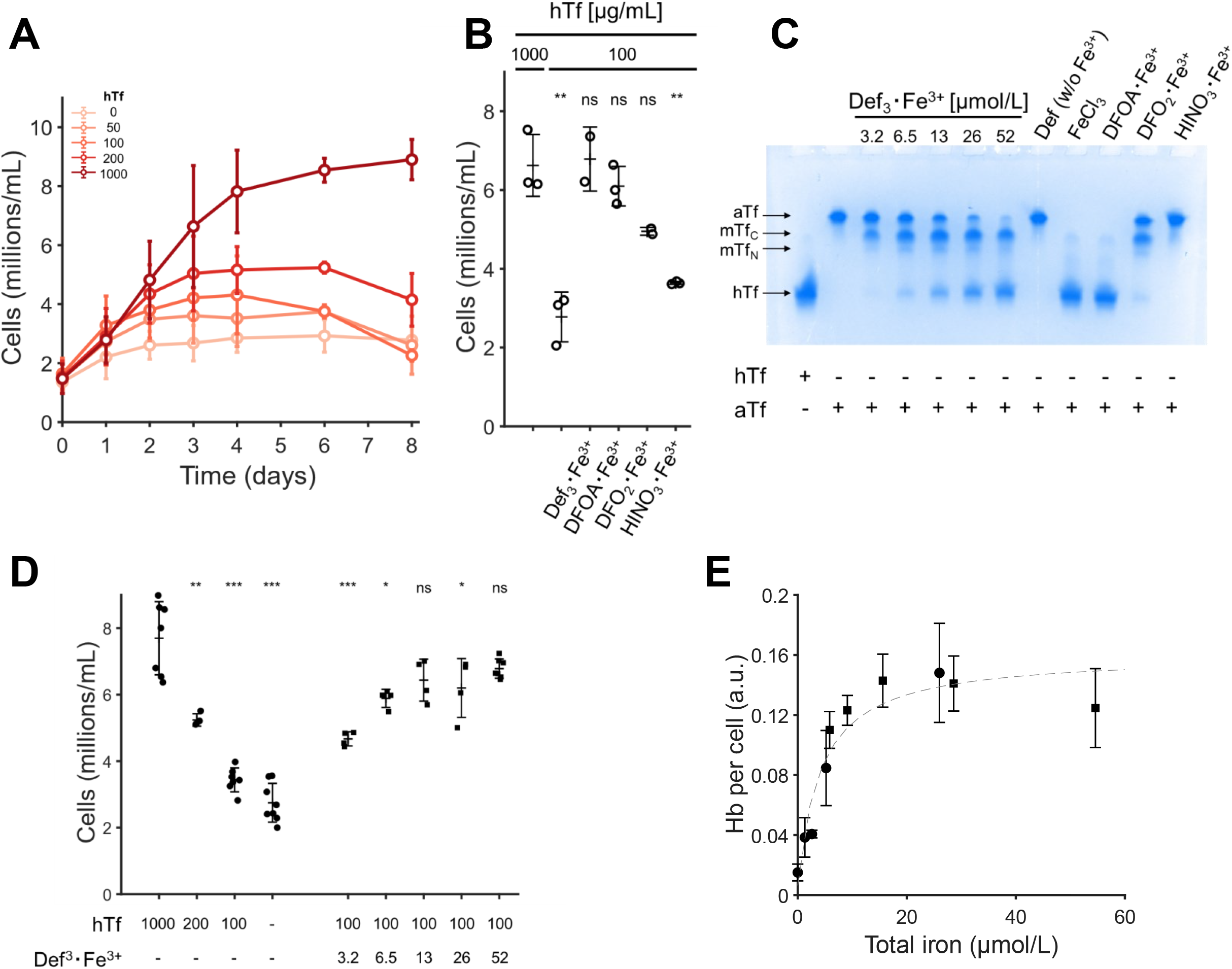
Deferiprone supplementation restores efficient differentiation of erythroblasts in transferrin-limited medium. Erythroblasts were expanded from PBMCs for 10-12 days, and subsequently seeded in differentiation medium at a starting cell concentration of 1.5 – 2.0 million cells/mL. (**A)** Cells were seeded with decreasing holotransferrin concentrations (1000, 200, 100, 50 and 0μg/mL). Cell number was determined at indicated days. **(B)** Cell concentration of at day 7 days of differentiation in medium with low hTf (100μg/mL), with or without 52μmol/L Fe^3+^ associated with one of four iron-loaded chelators (deferiprone = Def; deferoxamine = DFOA; deferasirox = DFO; hinokitiol = HINO). **(C)** 1000μg/mL apotransferrin was incubated for 16h with iron-loaded chelators (52μmol/L of chelated Fe^3^+, unless indicated). All samples were incubated overnight in PBS at 37°C (pH = 7.4). All samples were mixed 1:1 with Novex^™^ TBE-Urea gel sample buffer, loaded in a Novex^™^ 6% TBE-urea gel (3.0 μg of Tf per well; Invitrogen; Waltham, MA), and run for 2.25h at 150 V. Gels were stained with Coomassie. **(D)** Cell concentration after 6 days of differentiation in medium with hTf (values in μg/mL) and Def_3_·Fe^3+^ (values in μmol/L) as indicated. **(E)** Hemoglobin content per cell after 4 days of differentiation at increasing iron concentration in cells cultured using hTf as sole iron source (●), or Def_3_·Fe^3+^ plus 100μg/mL hTf (■) A hyperbolic dose-response curve is fitted on the data (Hb = Hb_max_ × [Fe^3+^] × (EC_50_ + [Fe^3+^])^-1^). All data is displayed as mean ± SD (error bars; n≥3). Significance is shown for the comparison with 1000μg/mL hTf (unpaired two-tailed two-sample Student’s *t*-test; ns for not significant differences, * for P<0.05, ** for P<0.01, *** for P<0.001).

#### Supplementation with iron-loaded chelators allows to reduce hTf concentrations

Iron-loaded holotransferrin is internalized upon binding to the transferrin receptor (TfR). After releasing Fe^3+^, iron-depleted transferrin (apotransferrin; aTf) is recycled back to the cell membrane together with the TfR. We tested whether supplementation of culture medium with an iron-loaded chelator can sustain differentiation of erythroblasts at low hTf concentrations. Four chelators loaded with iron (52μmol/L chelated Fe^3+^) were added to medium with a growth-limiting hTf concentration (100μg/mL). Supplementation with iron-loaded deferiprone (Def_3_·Fe^3+^) or deferoxamine (DFOA·Fe^3+^) enabled proliferation similar to 1000μg/mL of hTf, while both deferasirox (DFO_2_·Fe^3+^) and hinokitiol (HINO_3_·Fe^3+^) showed limited growth recovery (Figure 1B).

To test the ability of these chelators to reload apotransferrin, 1000μg/mL aTf was incubated with 52μmol/L of chelated iron at 37°C overnight and subjected to urea-PAGE gel electrophoresis. The mobility of aTf is lower compared to hTf, while the monoferric transferrin forms (mTf_C_, mTf_N_) have an intermediate migration speed. Deferasirox and hinokitiol showed a low Tf saturation level, while both deferiprone and deferoxamine could reload most aTf (Figure 1C). Soluble iron, added as FeCl_3_, was also able to reload the majority of aTf, with minor amounts of both mTf forms present. Although deferiprone resulted in more mTf compared to deferoxamine, the former was favored for stability^34^ and cost-effectiveness.

Decreasing deferiprone concentrations reduced the final saturation level of Tf (Figure 1C). To further test the effect of iron-loaded deferiprone supplementation (Def_3_·Fe^3+^) on iron-limited erythroid differentiation, erythroblasts were seeded in differentiation medium supplemented with 100μg/mL hTf supplemented and Def_3_·Fe^3+^ at concentrations between 3.2 and 52μmol/L (equivalent to 125 and 2000μg/mL hTf, respectively). Def_3_·Fe^3+^ improved cell growth dependent on the Def_3_·Fe^3+^ concentration (Figure 1D).

In differentiation cultures, most of the hemoglobin is synthesized in the first 2-4 days of culture, and hemoglobin content per cell is strongly dependent on the hTf concentration in the medium (Supplemental Figure S1B). At day 4, the hemoglobin content of cultured cells showed saturation kinetics (hyperbolic) as a function of the media iron content. Of note, cells cultured in absence of hTf had a 10-fold decrease in hemoglobin (Hb) mass per cell compared to the highest hTf dose tested (Figure 1E). This dose-response behavior was also observed when using 100μg/mL hTf with increasing deferiprone concentrations, with a maximal response using >13μmol/L Def_3_·Fe^3+^. The same iron-dependent response was seen for hemoglobin accumulation (Hb intracellular concentration; Supplemental Figure S1C-D), with no difference observed between Def_3_·Fe^3+^ concentrations >13μmol/L and medium containing 1000μg/mL hTf.

The data suggests that Def_3_·Fe^3+^ supplementation can reload apotransferrin generated during culture. Chemical equilibria simulations considering the affinities of ferric ions to the multiple species of deferiprone and transferrin support the observation that Def_3_·Fe^3+^ can reload iron-depleted transferrin, even by assuming the leaching of Fe3+ as the sole mechanism of iron transfer between the two chelators (Supplemental Figure S2).

### Iron-loaded deferiprone can fully replace holotransferrin in differentiation

Due to the lipophilic nature of deferiprone and its small size, it is expected to be cell permeable.^35^ We tested whether the larger Def_3_·Fe^3+^ complex can support erythroid differentiation in absence of transferrin. Expanded erythroblast cultures (day 12) were reseeded at 1.5×10^6^ cells/mL in differentiation medium with increasing concentrations of Def_3_·Fe^3+^ as sole iron source. At day 6 of differentiation, 3 times higher cell numbers were obtained using >13μmol/L Def_3_·Fe^3+^ compared with no hTf and no Def_3_·Fe^3+^ cultures (Figure 2A). Similar dose-response curves were obtained for total hemoglobin per cell (Figure 2B). The addition of increasing hTf concentrations to optimal concentrations of Def_3_·Fe^3+^ (52μmol/L) did not further increase cell number or hemoglobin content (Figure 2B-C). Interestingly, the combination of high levels Def_3_·Fe^3+^ together with a high concentration hTf did not lower cell numbers, suggesting that Def_3_·Fe^3+^ is not toxic to erythroblasts.

**Figure 2.**
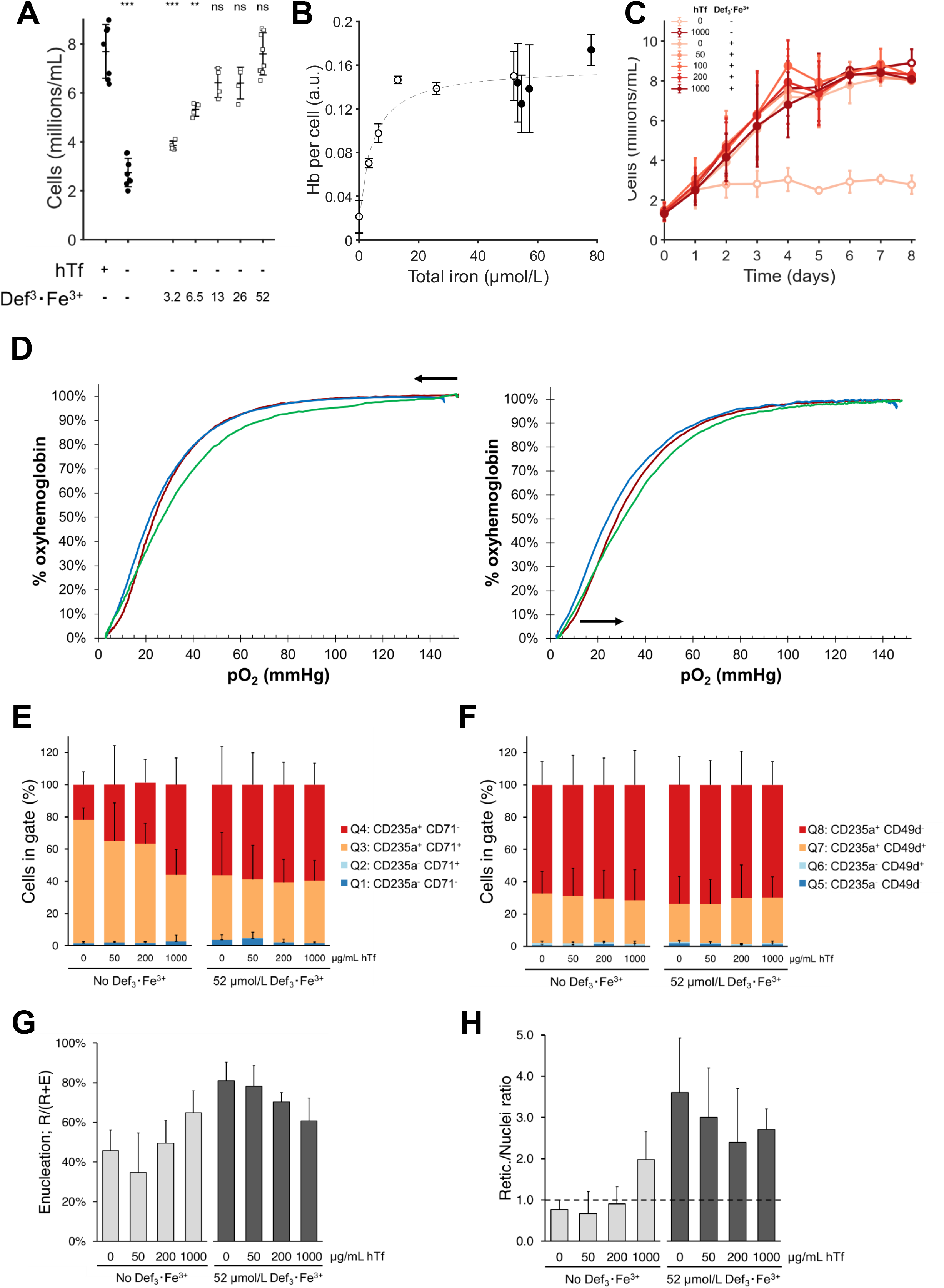
Deferiprone can fully replace holotransferrin in erythroblast differentiation cultures. Erythroblast were expanded from PBMCs for 10-12 days, and subsequently seeded in differentiation medium. **(A)** Cell concentration after 6 days of differentiation in medium without hTf supplemented with Def_3_·Fe^3+^ (concentrations indicated in μmol/L). **(B)** Hemoglobin content of cells after 4 days in differentiation in medium using Def_3_·Fe^3+^ as sole iron source (○), or with varying hTf concentrations (1000, 500, 200, 100, 50μg/mL) in the presence (●) of 52μmol/L Def_3_·Fe^3+^. **(C)** Cell concentration of erythroblasts seeded in medium with or without 52 μmol/L Def_3_·Fe^3+^ (filled and empty symbols, respectively), plus a decreasing concentration of hTf (values in μg/mL). **(D)** Oxygen dissociation (left) and association (right) curves measured by a HEMOX analyzer for peripheral blood erythrocytes (red), reticulocytes cultured using either 1000μg/mL hTf (blue) or 52μmol/L Def_3_·Fe^3+^ (green) as sole iron source. P50 values were calculated as the oxygen partial pressure that leads to a 50% saturation of hemoglobin. **(E-H)** Cells, differentiated for 10 days in presence of hTf and/or Def_3_·Fe^3+^ as indicated, were stained with CD235 plus CD71 **(E)**, CD235a plus CD49d **(F)**, or DRAQ5 (cell permeable DNA stain; **G, H**). Relative cell numbers per quadrant were calculated (gating strategy available in Supplemental Figure S3A). **(G)** Enucleation percentage of erythroid cells, and **(H)** ratio of reticulocytes versus pyrenocytes (extruded nuclei) were calculated from the forward scatter and DRAQ5 staining. A retic./nuclei ratio > 1 means more reticulocytes than nuclei. All data is displayed as mean ± SD (n≥3). Significance is shown for the comparison with the 1000μg/mL hTf condition (unpaired two-tailed two-sample Student’s *t*-test; ns for not significant differences, * for P<0.05, ** for P<0.01, *** for P<0.001).

Oxygen association and dissociation are crucial for RBC function. Reticulocytes obtained from differentiation cultures using 52μmol/L Def_3_·Fe^3+^ as sole iron source had similar hemoglobin oxygen dissociation and association curves compared to peripheral blood RBCs and reticulocytes obtained using hTf, with a slightly higher P_50_ (blood = 26.8 mmHg, hTf = 24.2 mmHg, deferiprone = 28.0 mmHg; calculated from oxygenation profile; Figure 2D).

Flow cytometry studies were performed to evaluate the effect of hTf and Def_3_·Fe^3+^ on the erythroblast maturation level. During differentiation erythroblast acquire CD235 (Glycophorin A), and eventually lose CD71 (TfR1) expression. Expression of CD71 may not be a reliable differentiation marker when evaluating the effect of hTf and Def_3_·Fe^3+^ supplementation as it is upregulated at low iron availability.^36^ At day 10 of differentiation, cells had a high expression of CD235 under all tested conditions (gating strategy in Supplemental Figure S3A). In contrast, CD71 expression levels decreased with increasing iron availability, independent of whether iron is presented as hTf or Def_3_·Fe^3+^ (Figure 2E; Supplemental Figure S3B). CD49d (integrin alpha 4) is an early differentiation marker^37^ expressed independent of iron metabolism regulation. Regardless of the hTf or Def_3_·Fe^3+^ concentration, 30%-40% of all cells had a CD235^+^ CD49d^+^ phenotype, suggesting that deferiprone supplementation did not delay erythroid differentiation (Figure 2F).

The availability of chelated iron has been reported to affect erythroblast enucleation.^38^ Enucleation efficiency of the cultures was evaluated using the nuclear stain DRAQ5 and cell size by flow cytometry (Supplemental Figure S3A). The ratio between enucleated (R) and total cells (R+E) indicates the enucleation ratio. High hTf concentrations led to a slightly higher enucleation efficiency (65%) compared with suboptimal hTf concentrations (0μg/mL hTf: 46%; 50μg/mL hTf: 35%; 200μg/mL hTf: 50%) (Figure 2G). The effect of iron limitation was stronger on reticulocytes and nuclei stability, with a 2-to 3-fold decrease in the reticulocyte to nuclei ratio (R/N) compared with cultures with 1000μg/mL hTf. Addition of 52μmol/L Def_3_·Fe^3+^ increased terminal enucleation and recovered reticulocyte/nuclei ratio (Figure 2H). Collectively, these results suggest that Def_3_·Fe^3+^ can fully replace hTf in differentiation cultures, maintaining high cell yields and enucleation efficiency, hemoglobin content and oxygen carrying capacity.

### Def_3_·Fe^3+^ supplementation prevents molecular responses to iron deficiency

Next, we studied whether iron supplementation by Def_3_·Fe^3+^ and hTf similarly controlled cellular iron metabolism in erythroblasts. The IRP1/2-dependent expression of ferritin and TfR, and HRI-dependent phosphorylation of eIF2 was measured during the first 2 days of differentiation in presence of high and low levels of hTf or Def_3_·Fe^3+^ (Figure 3A). The expression of TfR was upregulated in presence of apotransterrin, a condition in which all available iron is quenched. This is in accordance with IRP stabilizing TfR mRNA in iron-limiting conditions. Low iron concentrations did not significantly affect TfR expression, possibly due to the stability and recycling of the TfR (quantification: Figure 3B).

**Figure 3.**
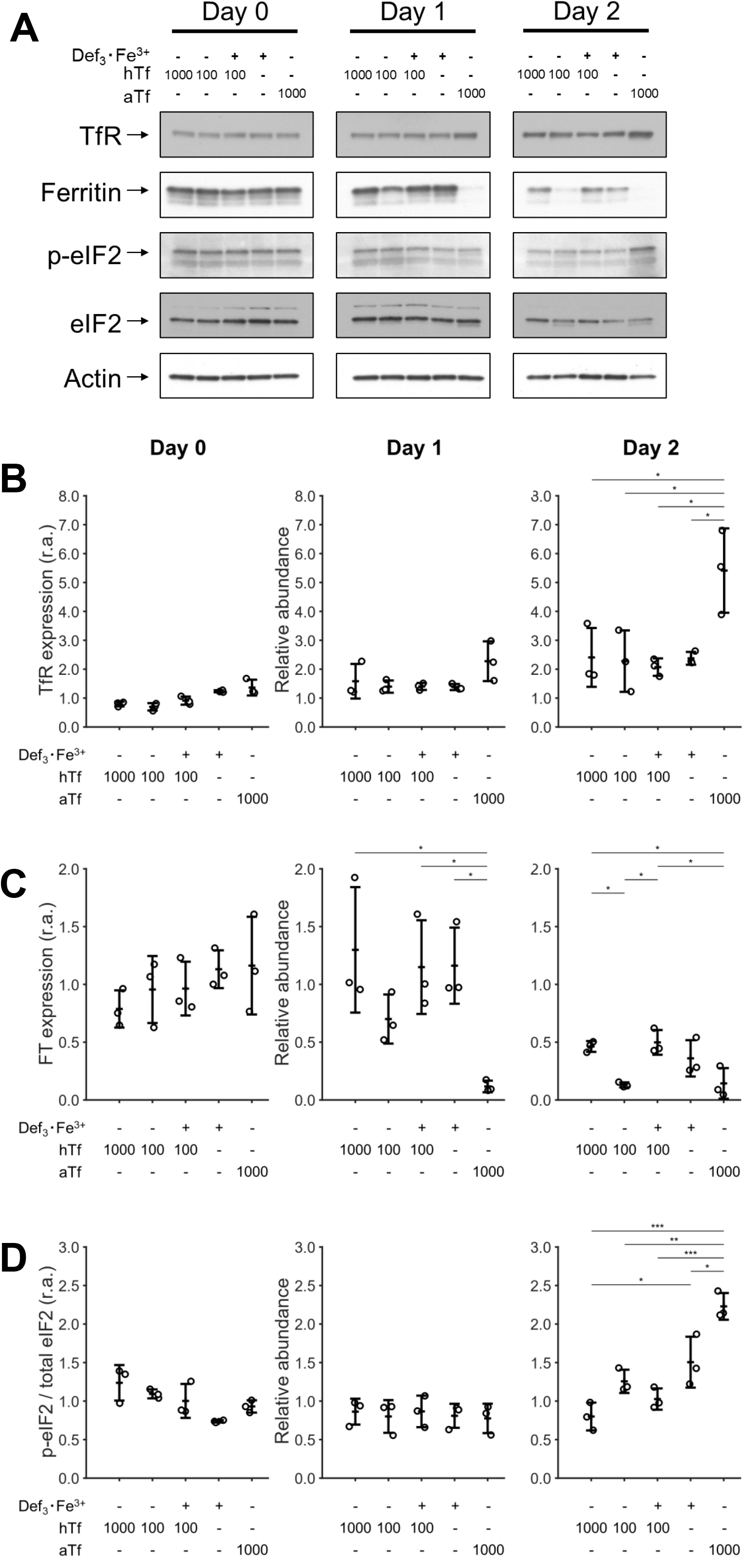
Recovery of iron regulation metabolism using deferiprone in differentiation. Erythroblasts were expanded from PBMCs for 10-12 days, and subsequently seeded in differentiation medium supplemented with hTf or aTf (values μg/mL) or Def_3_·Fe^3+^ (52μmol/L). Cells were harvested at seeding (Day 0) and after 1 and 2 days of culture. **(A)** Western blots of one representative donor showing the expression level of proteins involved in iron metabolism regulation. Actin was used as housekeeping protein for relative quantification of protein abundance. **(B-C)** Relative abundance (r.a.) of transferrin receptor (TfR) and ferritin. Relative protein levels were calculated using actin signal, followed by normalization using the average of each donor at the start of the culture (day 0). **(D)** Level of the phosphorylated form of eIF2 (p-eIF2) relative to the total levels of eIF2. In order to compare different experiments, the relative p-eIF2/eIF2 ratio was normalized using the average of each donor at the start of the culture (day 0). Data is displayed as mean ± SD (error bars; n=3). Significance is shown for the comparison with 1000μg/mL hTf (unpaired two-tailed two-sample Student’s *t*-test with Helm-Bonferrin correction for multiple comparisons; * for P<0.05, ** for P<0.01, *** for P<0.001).

Ferritin synthesis, in contrast, is inhibited upon IRP binding to ferritin mRNA.^12^ Ferritin levels remained high in presence of high total iron concentrations (1000μg/mL hTf, or 52μmol/L Def_3_·Fe^3+^), but were reduced at low or absent iron levels (100μg/mL hTf, or apotransferrin) (Figure 3C). Finally, high levels of p-eIF2 were observed in cultures with iron limitation (Figure 3D; day 2). Def_3_·Fe^3+^ supplementation restored the low phosphorylation levels of eIF2 detected in cultures with high hTf concentration. Together, our results suggest that iron delivery to erythroblasts by Def_3_·Fe^3+^ or through hTf is fully interchangeable at the molecular level.

### Medium containing iron-loaded deferiprone can sustain expansion of erythroblasts

Iron requirements are especially high during erythroblast differentiation when hemoglobin is synthesized. However, cells do require iron as constituent of essential metalloproteins, such as cytochromes and some peroxidases.^39^ We tested if deferiprone could also replace hTf during the first phase of our culture protocol, when hematopoietic stem and progenitor cells commit to the erythroid lineage, and subsequently when committed erythroblasts proliferate but hemoglobin production remain low. Erythroid cultures were established from peripheral blood mononuclear cells (PBMCs) in medium supplemented with the standard hTf concentration for culturing of non-hemoglobinized cells (300μg/mL hTf), or with a 10x lower hTf concentration, in presence or absence of Def_3_·Fe^3+^ (52μmol/L). The presence of hTf (300μg/mL) was required to obtain a culture of CD71^+^ cells from PBMCs (Figure 4A; Supplemental Figure S3A-B), while deferiprone alone could not replace hTf to establish an erythroid culture. At a lower hTf concentration (30μg/mL) cultures yielded less CD71^+^ erythroblasts, which was unaffected if deferiprone was supplemented. As shown previously, CD71^+^ cells cultured in absence of hTf expressed CD71 at increased levels. In these early stages, deferiprone did not reduce CD71 levels to those observed with hTf (Supplemental Figure S3C). Upon prolonged expansion of committed erythroblasts Def_3_·Fe^3+^, alone or supplemented to low hTf levels, resulted in the same erythroblast growth rate as when 300μg/mL hTf was used (Figure 4B).

**Figure 4.**
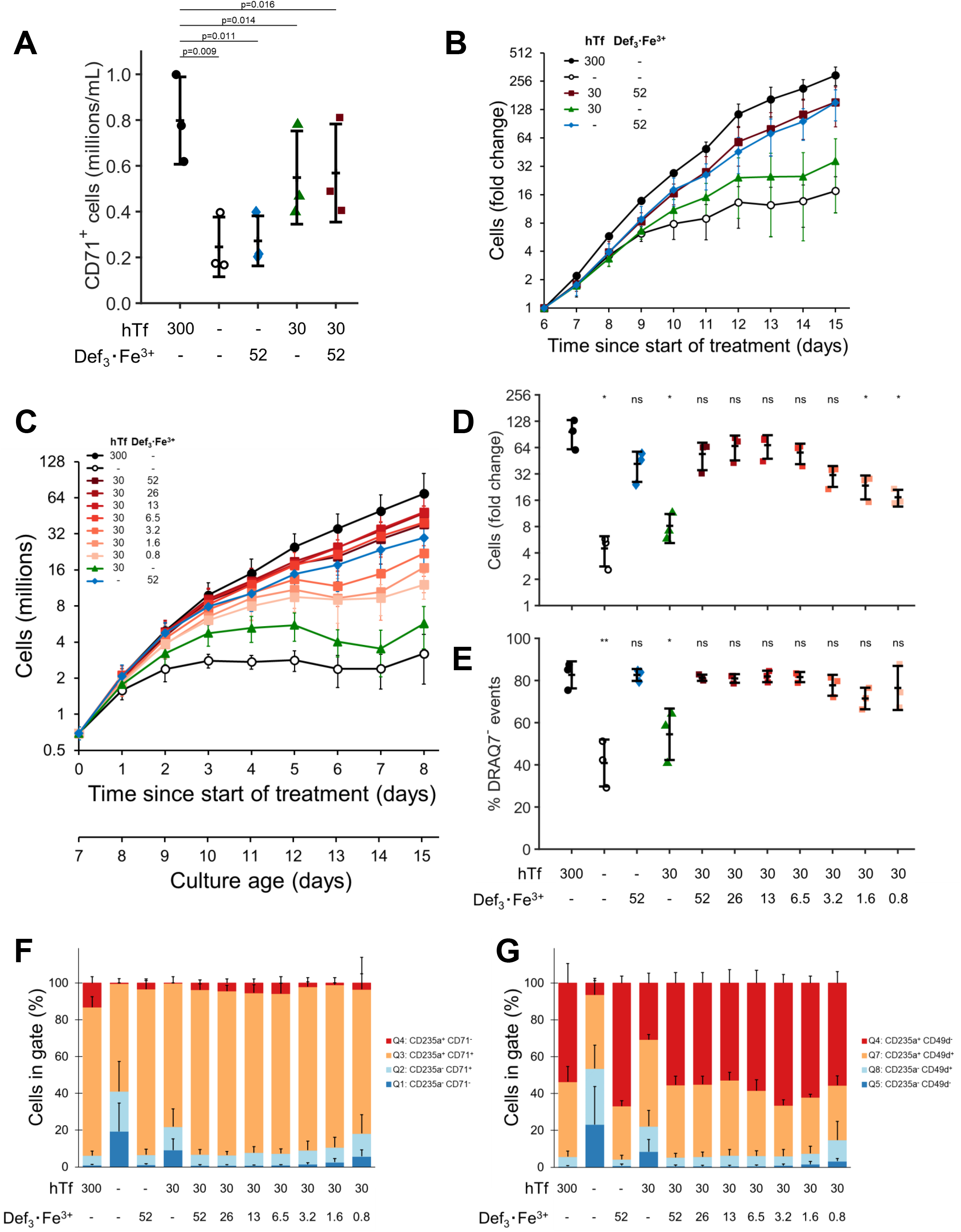
Deferiprone supplementations sustains erythroblast expansion. Adult PBMCs were isolated (day 0) and cultured in expansion medium at a starting cell concentration of 1 million cells/mL using hTf (values in μg/mL) in the presence or absence of Def_3_·Fe^3+^ (52μmol/L). **(A)** Total number of CD71^+^ cells was determined on day 6 after PBMC isolation (combining cell count and CD71+ frequency by flow cytometry). **(B)** Erythroblast cell concentration was monitored for additional 9 days of culture, with daily medium refreshment. Fold change in cell number was calculated using the measured number of erythroblasts at day 6. **(C-G)** Erythroblasts were expanded from PBMCs in expansion medium with hTf as iron source (300μg/mL) for 7 days, and subsequently reseeded in medium with varying hTf concentrations (300, 30, 0μg/mL) in the presence of Def_3_·Fe^3+^ (0.8 – 52μmol/L Def_3_·Fe^3+^). Cultures were maintained for 8 days (culture age = 15 days since PBMC isolation) during which daily medium refreshments were performed **(C)**. After 8 days of culture, cell number fold change was calculated relative to the start of treatment **(D)**. Cells were stained with DRAQ7 (cell impermeable DNA stain; **E**), CD235a plus CD71 **(F)**, and CD235a plus CD49d **(G)**. Relative cell numbers per quadrant were calculated (representative density plots available in Supplemental Figure 5). Data is displayed as mean ± SD (error bars; n=3). Significance is shown for the comparison with 300μg/mL hTf (paired two-tailed two-sample Student’s *t*-test; ns for not significant differences, * for P<0.05, ** for P<0.01, *** for P<0.001).

To determine the optimal Def_3_·Fe^3+^ concentration to sustain proerythroblast proliferation once erythroid cultures are established, PBMCs were first expanded for 6 days in medium with hTf, followed by culture in medium with a suboptimal hTf level supplemented with Def_3_·Fe^3+^ at concentrations between 0.8 and 52μmol/L (Figure 4C). Maximum recovery required at least 6.5μmol/L Def_3_·Fe^3+^ (Figure 4D). Lack of iron, or low levels of iron (30μg/mL hTf) increased the number of non-viable cells as detected by DRAQ7. Supplementation with Def_3_·Fe^3+^ recovered viability of the expansion cultures (Figure 4E). During the expansion phase of erythroblast cultures, the CD71^high^ erythroblast population slowly gains CD235 expression from day 10 onwards.^25^ A CD71^mid^ CD235^-^ subpopulation was observed under iron-limited conditions (0 and 30μg/mL hTf), and less prominently under low Def_3_·Fe^3+^ concentrations (1.6 and 0.8μmol/L) (Figure 4F; Supplemental Figure S5). The presence of these seemingly non-erythroid cells was also observed when following the expression of CD49d (Figure 4G). High levels of Def_3_·Fe^3+^ enhanced the gain of CD235 and loss of CD49d to levels observed with 300μg/mL hTf.

### Iron-loaded deferiprone can replace hTf for other myeloid cell lines

Next, we evaluated the potential of Def_3_·Fe^3+^ to fully replace holotransferrin in the myeloid cell lines MOLM13, NB4, EOL1, K562, HL-60, and ML-2. Cells were seeded in media containing no iron, 300μg/mL hTf or 52μmol/L Def_3_·Fe^3+^. All cell lines could be expanded in our serum-free medium supplemented with hTf (Figure 5A; Supplemental Figure S6). Lack of iron decreased the growth rate, with MOLM13 and NB4 cells showing a decrease of 4 orders of magnitude in cell numbers after 16 days of culture. EOL1 cultures seemed less sensitive to lack of hTf, but proliferation in presence of hTf was also low. Def_3_·Fe^3+^ recovered growth to levels comparable with only-hTf conditions. As observed in *ex vivo* erythroblast expansion and differentiation cultures, lack of hTf led to an upregulation of CD71 expression in all cell lines (Figure 5B). Def_3_·Fe^3+^ supplementation restored CD71 expression levels comparable to that in hTf-only cultures.

**Figure 5.**
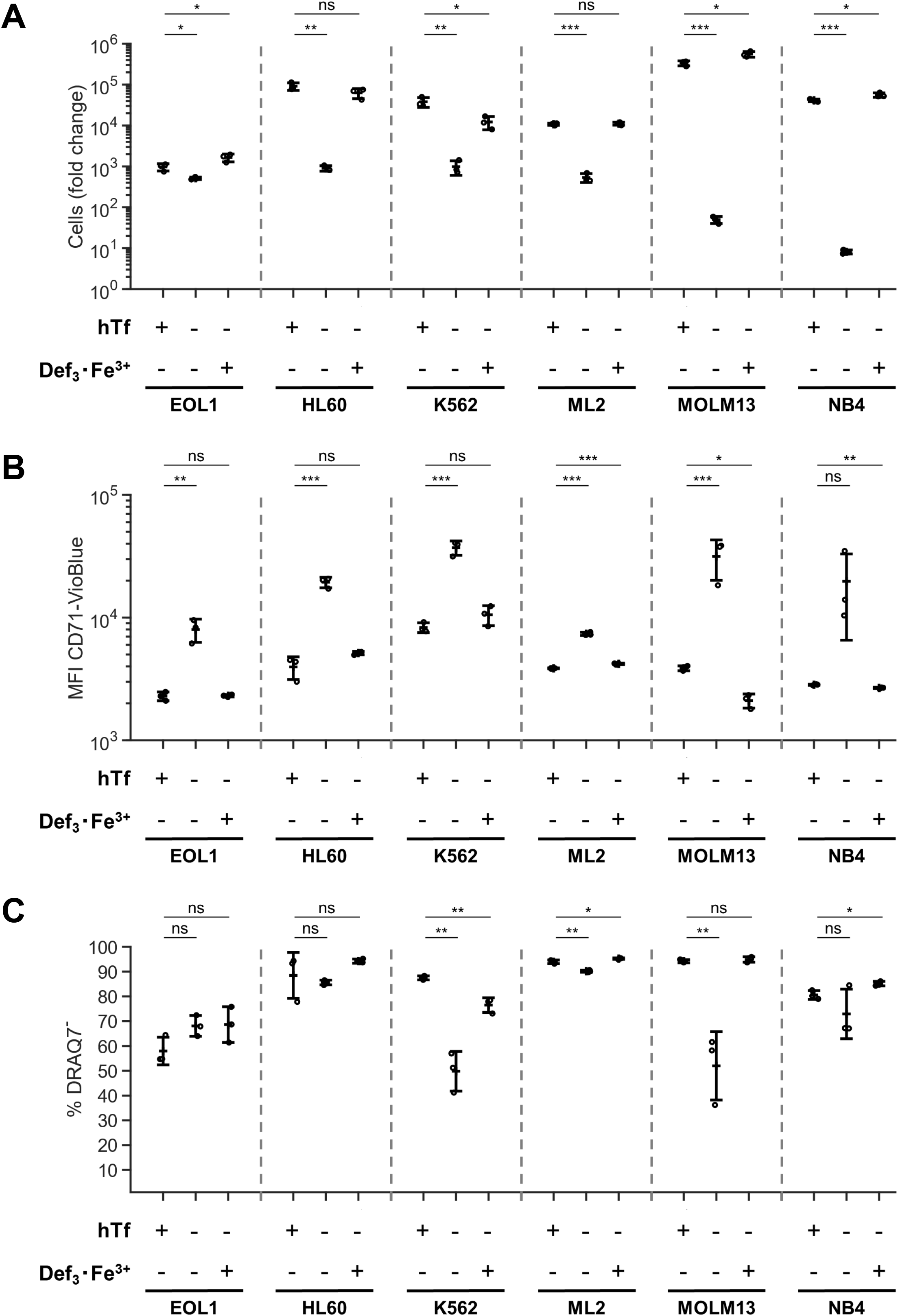
Deferiprone sustains expansion of selected myeloid cell lines. Myeloid cell lines were cultured in Cellquin medium supplemented or not with 300μg/mL hTf or 52μmol/L Def_3_·Fe^3+^ for 16 days. **(A)** Total cell number fold change relative to day 0. **(B)** Expression level of the transferrin receptor (CD71) determined by flow cytometry. MFI for corresponding isotypes is between 1×10^2^ - 3×10^2^. **(C)** Cells were stained with DRAQ7 (cell impermeable DNA stain) to quantify the viability of the cultures. Data is displayed as mean ± SD (error bars; individual datapoints are indicated; n=3). Significance is shown for the comparison with 300 μg/mL hTf (unpaired two-tailed two-sample Student’s *t*-test; ns for not significant differences, * for P<0.05, ** for P<0.01, *** for P<0.001).

## Discussion

Hemoglobin production requires a high uptake of iron in differentiating erythroblasts, which is supplied as hTf. We evaluated whether iron-loaded chelators can be a low-cost alternative for hTf in *ex vivo* erythroid cultures. Our data show that deferiprone, a bidentate α-ketohydroxypyridine iron-specific chelator, could completely replace holotransferrin to produce large numbers of enucleated hemoglobinized cultured RBCs at concentrations as low as 26μmol/L (Def_3_·Fe^3+^). Hemoglobin content and oxyhemoglobin dissociation dynamics of cultured RBCs (cRBCs) derived with Def_3_·Fe^3+^ or with hTf as sole iron source were similar.

### Deferiprone: a cell permeable carrier of iron

We show that hemoglobin is expressed at similar levels, and is equally functional in reticulocytes cultured in presence of hTf or Def_3_·Fe^3+^. The ability of deferiprone to permeate and access labile iron pools in the cytosol, mitochondria and nuclei may be due to its low molecular weight (139.1g/mol) and mild lipophilic nature (D_7.4_ = 0.17), as previously reported.^40,41^ Intracellular mobilization of iron by deferiprone, and the availability of deferiprone-bound iron for hemoglobin synthesis has also been described.^24^ Other iron-loaded chelators such as DFOA·Fe^3+^ can also function as substrate for heme synthesis instead of holotransferrin.^42^ Excessive iron availability, however, may also cause oxidative damage.^43,44^ High levels of Def_3_·Fe^3+^ (26 – 52μmol/L) did not have a detrimental effect on cell yield or enucleation efficiency, indicating that the iron-binding affinity of deferiprone is just right to prevent toxicity of free iron, while simultaneously releasing iron to cellular proteins. Our data suggests that Def_3_·Fe^3+^ acts independent of Tf and the TfR, but only deletion of TfR1/TfR2 will confirm this. The availability of iron upon Def_3_·Fe^3+^ treatment on cultured erythroblasts increased ferritin expression, indicating that Def_3_·Fe^3+^ can release iron to ferritin. We propose to evaluate Def_3_·Fe^3+^ intracellular trafficking by radiolabeled iron tracing in cells at the different stages of our culture protocol, to determine how iron carried by deferiprone is mobilized and incorporated into the pathways of iron metabolism in early and late erythroid cells.

Interestingly, phosphorylation levels of eIF2α were higher in cells cultured with Def_3_·Fe^3+^. eIF2α can be phosphorylated by the heme-regulated eIF2α-kinase (HRI), which is activated under heme-limited conditions in erythroid cells.^13^ As hemoglobin levels were maintained in Def_3_·Fe^3+^-treated cells, it is possible that the observed increase in p-eiF2α is due to other mechanisms, such as an increase in intracellular oxidative stress levels due to iron overload.

### Non-committed erythroid progenitors display an inability to use Def_3_·Fe^3+^

Def_3_·Fe^3+^ in absence of hTf supported erythroblast expansion during the second stage of our culture system. However, Def_3_·Fe^3+^ was unable to support the establishment of erythroblast-enriched cultures from PBMCs. The addition of suboptimal concentrations of hTf to these cultures allows for limited outgrowth of erythroblasts, which was not altered by the presence of Def_3_·Fe^3+^.This suggests that toxicity by iron overload is not causing the lower yield. In contrast, once cells were committed to the erythroid lineage, Def_3_·Fe^3+^ could substitute hTf. We hypothesize that in the first days of culture, precursor cells not yet committed to the erythroid lineage may lack the molecular machinery to incorporate iron ions provided by Def_3_·Fe^3+^. It becomes relevant to understand how Def_3_·Fe^3+^ complexes are processed by the cells and their interaction with intracellular iron pools, as the data suggests a differential ability between hematopoietic progenitors and erythroblasts to utilize this source of iron. Of note, these early cultures, however, are small scale and the use of hTf is not yet an important cost factor. Thus, cultures could be started in presence of hTf, and be continued in medium supplemented with Def_3_·Fe^3+^ once an erythroblast-enriched population is established.

### Outlook

Holotransferrin is the main iron source in mammalian cell culture medium, and is typically included in culture media by supplementation with serum. However, serum-containing media have an unclear chemical composition, typically show batch-to-batch variation, and have a risk of contamination; for example, by viruses. Defined serum-free formulations have been developed, and require purified hTf.^45^ Use of purified Tf from plasma, or of recombinant Tf from rice or yeast, can potentially introduce contaminants. Def_3_·Fe^3+^, however, is chemically produced, which can be an advantage with respect to purity in the production of chemically-defined protein-free medium.

Migliaccio et al.^5^ proposed the first serum-free medium formulation for erythroid culture, and showed that transferrin levels have a direct impact on the yields of erythroid progenitor cells. Erythroblast expansion and differentiation can be achieved using 300 and 1000μg/mL hTf, respectively.^25^ The results of our study indicate that although transferrin is currently essential for the establishment of erythroid cultures from PBMCs, it can be fully replaced by Def_3_·Fe^3+^ during erythroblast expansion, differentiation and maturation. As transferrin represents 40-50% of the total medium cost in erythroid differentiation cultures, replacing it by this chelator would result in a decrease of media cost of $200-250 per liter. Although hTf concentrations are significantly lower in culture media for other primary cells and for cell lines typically used in biopharmaceutical processes (e.g. hybridoma, CHO, Vero; ranging between 5 and 100μg/mL), its replacement with Def_3_·Fe^3+^ may be a significant cost-saver at an industrial production scale.

## Acknowledgements

We thank Paul Kaijen and Jan Voorberg (Molecular Hematology, Sanquin Research) for technical help and advice on iron loading of transferrin. This work was supported by the ZonMW TAS program (project 116003004), by the Landsteiner Foundation for Bloodtransfusion Research (LSBR project 1239), and by Sanquin Blood Supply grants PPOC17-28 and PPOC119-14. E.M.P was supported by an Erasmus Fellowship (project WORK4ALL 2018-1-PT01-KA103-046800).

## Authorship

### Contribution

J.S.G.M., N.Y., E.M.P., and A.A.V. performed the experiments; J.S.G.M., and M.v.L. designed the experiments, analyzed the data, and wrote the manuscript; A.W., and E.v.d.A. contributed to the experiment design and writing of the manuscript. All authors critically revised the manuscript.

