## Supplemental Figures 1-6 for "Iron-loaded deferiprone can support full hemoglobinization of cultured red blood cells in the absence of transferrin"

**Supplemental Figure 1. Intracellular hemoglobin concentration of differentiating erythroblasts using holotransferrin and iron-loaded deferiprone as iron source.** Erythroblasts were expanded from PBMCs for 10-12 days, and subsequently seeded in differentiation medium at a starting cell concentration of 1.5 – 2.0 million cells. **(A)** Cells were seeded with decreasing holotransferrin concentrations (1000, 200, 100, 50 and 0µg/mL). Evolution of mean cell diameter is depicted. **(B)** Hemoglobin cell content (i.e. mass of hemoglobin per cell; displayed as arbitrary units = absorbance at 420 nm per million cells) was determined at indicated days. The effect of deferiprone supplementation with sub-optimal hTf concentrations on erythroblast cell volume **(C)** and hemoglobin content **(D)** in differentiation cultures was also evaluated. For this, erythroblast expanded from PBMCs were seeded in medium using hTf as sole iron source (dark grey; ●), or in hTf-limited conditions (100µg/mL hTf) in the presence of $Def_3 \cdot Fe^{3+}$ in concentrations ranging between 3.2 and 52µmol/L (light grey; ■). Intracellular hemoglobin concentration was calculated using the cell volume and Hb per cell values (day 4 of treatment), and is depicted in arbitrary units (a.u.) of absorbance per fL of cell volume. Mean ± SD (error bars; n≥3). Significance is shown for the comparison with 1000µg/mL hTf (unpaired two-tailed two-sample Student's *t*-test, $^*p<0.05$; n≥3).

**Supplemental Figure 2. Modeling of iron saturation in transferrin and deferiprone solutions.** **(A)** Iron shuttling model proposed for the reloading of apotransferrin in culture using iron-loaded deferiprone. Leaching of $Fe^{3+}$ ions from deferiprone or transferrin was the only mechanism considered for the transfer of iron between the two chelators. **(B)** Concentrations of transferrin and deferiprone species were calculated assuming equilibria at culture conditions (pH = 7.4, 3.5mmol/L $HCO_3^-$). For transferrin, the association equilibrium constants of $7.0\times10^{22}$ L/mol and $3.6\times10^{21}$ L/mol for the binding of the first and second $Fe^{3+}$ ion were assumed, respectively.[46] No difference in the association of iron to the N- and C-lobes of transferrin was considered. For deferiprone, the global stability constants (log β) for the $Def\cdot Fe^{3+}$, $Def_2\cdot Fe^{3+}$ and $Def_3\cdot Fe^{3+}$ complexes were assumed to be 15.01, 27.30 and 37.43, respectively.[22] A constant average iron uptake rate of $1.7\times10^{-7}$ mol $Fe^{3+}$/L·h was assumed, corresponding to the production of $10\times10^{6}$ hemoglobinized cells per mL of culture (1 cell = 300 million Hb molecules) in 4 days. Calculation of transferrin and deferiprone concentrations for each timepoint was performed solving the system of nonlinear chemical equilibrium equations with MATLAB ver. R2019b. **(C)** Calculated time profiles for the concentrations of the different transferrin and deferiprone species for cultures with either 1000 or 100µg/mL of hTf, in presence or absence of $Def\cdot Fe^{3+}$. Under low hTf conditions, supplementation with $Def\cdot Fe^{3+}$ leads to a delay on the depletion of hTf.

**Supplemental Figure 3. Expression of erythroid cell surface markers in differentiation cultures supplemented with deferiprone.** Erythroblasts were differentiated for 10 days in medium at different hTf concentrations, in the presence or absence of $Def_3 \cdot Fe^{3+}$ (52µmol/L). **(A)** Gating strategy to evaluate the differentiation level of cultured erythroblasts. Cells were gated (FSC/SSC), followed by gating of single cells (FSC-A/FSC-H). Cells are depicted in a CD235a/CD71 or a CD235a/CD49d dot plot. To evaluate enucleation levels, erythroblasts ($DRAQ5^{+}$ $FSC^{high}$), pyrenocytes (extruded nuclei; $DRAQ5^{+}$ $FSC^{low}$) and reticulocytes ($DRAQ5^{-}$) were gated. **(B)** Representative density plots indicating the expression of the cell surface markers CD71, CD235 and CD49d after 10 days of culture.

**Supplemental Figure 4. Cell yields and expression of transferrin receptor TfR (CD71) on erythroid cultures stablished under deferiprone supplementation.** PBMCs were cultured in expansion medium without iron, or supplemented with holotransferrin (30µg/mL or 300µg/mL) and $Def_3 \cdot Fe^{3+}$ (52µmol/L), as indicated. After 6 days of culture, the typical time for the establishment of erythroid cultures, total cell concentration was measured **(A)**. Percentage of $CD71^{+}$ cells **(B)** and the mean fluorescence intensity of CD71 **(C)** were determined by flow cytometry. Cultures were kept for 9 days more, as shown in Figure 4B. Mean ± SD (error bar; n=3). Significance is shown for the comparison with the 300µg/mL hTf condition (paired two-tailed two-sample Student's *t*-test; ns for not significant differences, * for P<0.05, ** for P<0.01, *** for P<0.001).

**Supplemental Figure 5. Expression of erythroid cell surface markers in erythroblasts expanded in medium supplemented with deferiprone.** PBMCs were cultured in our original expansion medium (300µg/mL hTf) for 7 days until an erythroblast-enriched culture was obtained, followed by further culturing using 300, 30 or 0µg/mL hTf in the presence of $Def_3{\cdot}Fe^{3+}$ (0.8 – 52µmol/L $Def_3{\cdot}Fe^{3+}$). Representative density plots indicating the expression of the cell surface markers CD71, CD235 and CD49d curves after 8 days of treatment (culture age = 15 days since PBMC isolation).

**Supplemental Figure 6. Deferiprone sustains expansion of selected myeloid cell lines.** Six myeloid cell lines were cultured in serum-free expansion medium supplemented with 300µg/mL hTf or 52µmol/L $Def_3{\cdot}Fe^{3+}$ for 16 days. Cells were cultured under static conditions at a concentration of $0.3\times10^6$ cells/mL, with changing of medium every 2 days. Final cell number fold change relative to day 0 is depicted. Data is displayed as mean ± SD (error bar; n=3).

### Supplemental Figure 1

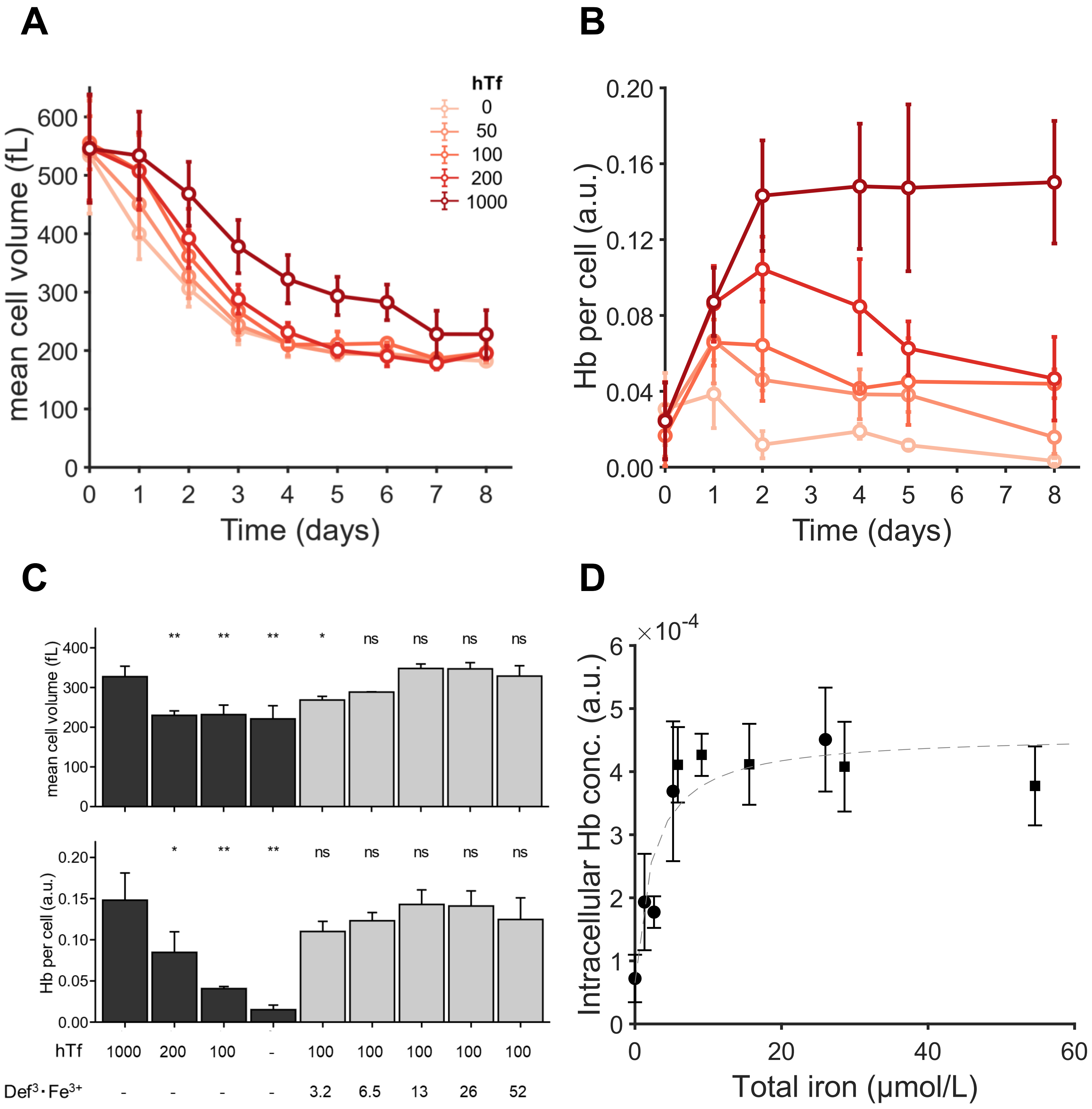

### Supplemental Figure 2

A

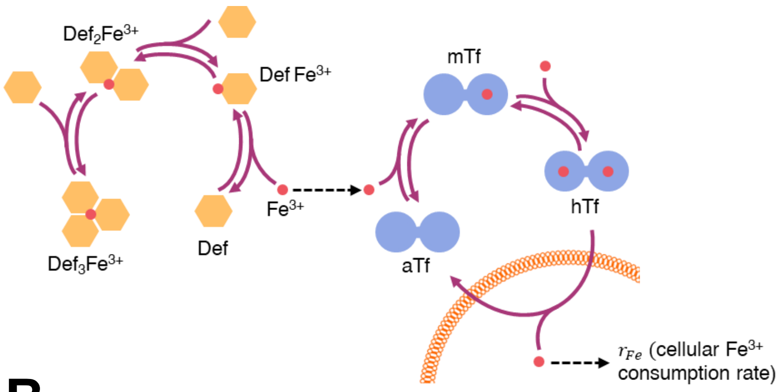


B

$$aTf + Fe^{3+} \underset{}{\overset{K_{Tf,1}}{\rightleftarrows}} mTf \qquad K_{Tf,1} = \frac{[mTf]}{[aTf][Fe^{3+}]} = 7.0 \times 10^{22}\ M^{-1}$$

$$mTf + Fe^{3+} \underset{}{\overset{K_{Tf,2}}{\rightleftarrows}} hTf \qquad K_{Tf,2} = \frac{[hTf]}{[mTf][Fe^{3+}]} = 3.6 \times 10^{21}\ M^{-1}$$

$$Def + Fe^{3+} \underset{}{\overset{\beta_{Def,1}}{\rightleftarrows}} Def \cdot Fe^{3+} \qquad \beta_{Def,1} = \frac{[Def \cdot Fe^{3+}]}{[Def][Fe^{3+}]} = 10^{15.01}\ M^{-1}$$

$$Def \cdot Fe^{3+} + Fe^{3+} \underset{}{\overset{\beta_{Def,2}/\beta_{Def,1}}{\rightleftarrows}} Def_2 \cdot Fe^{3+} \qquad \beta_{Def,2} = \frac{[Def_2 \cdot Fe^{3+}]}{[Def]^2[Fe^{3+}]} = 10^{27.30}\ M^{-2}$$

$$Def_2 \cdot Fe^{3+} + Fe^{3+} \underset{}{\overset{\beta_{Def,3}/\beta_{Def,2}}{\rightleftarrows}} Def_3 \cdot Fe^{3+} \qquad \beta_{Def,3} = \frac{[Def_3 \cdot Fe^{3+}]}{[Def]^3[Fe^{3+}]} = 10^{37.43}\ M^{-3}$$

C

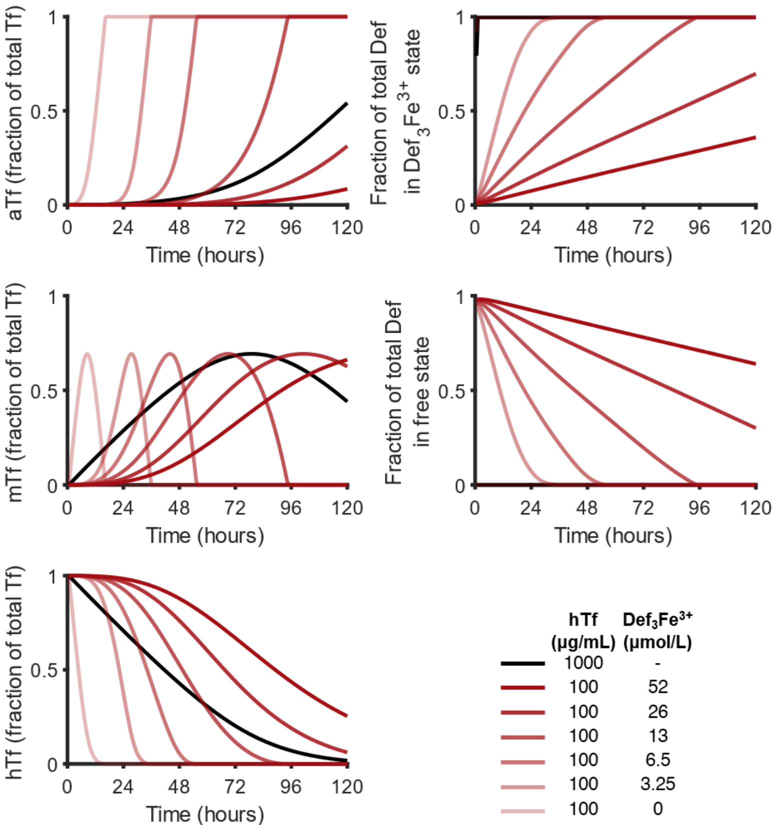

### Supplemental Figure 3

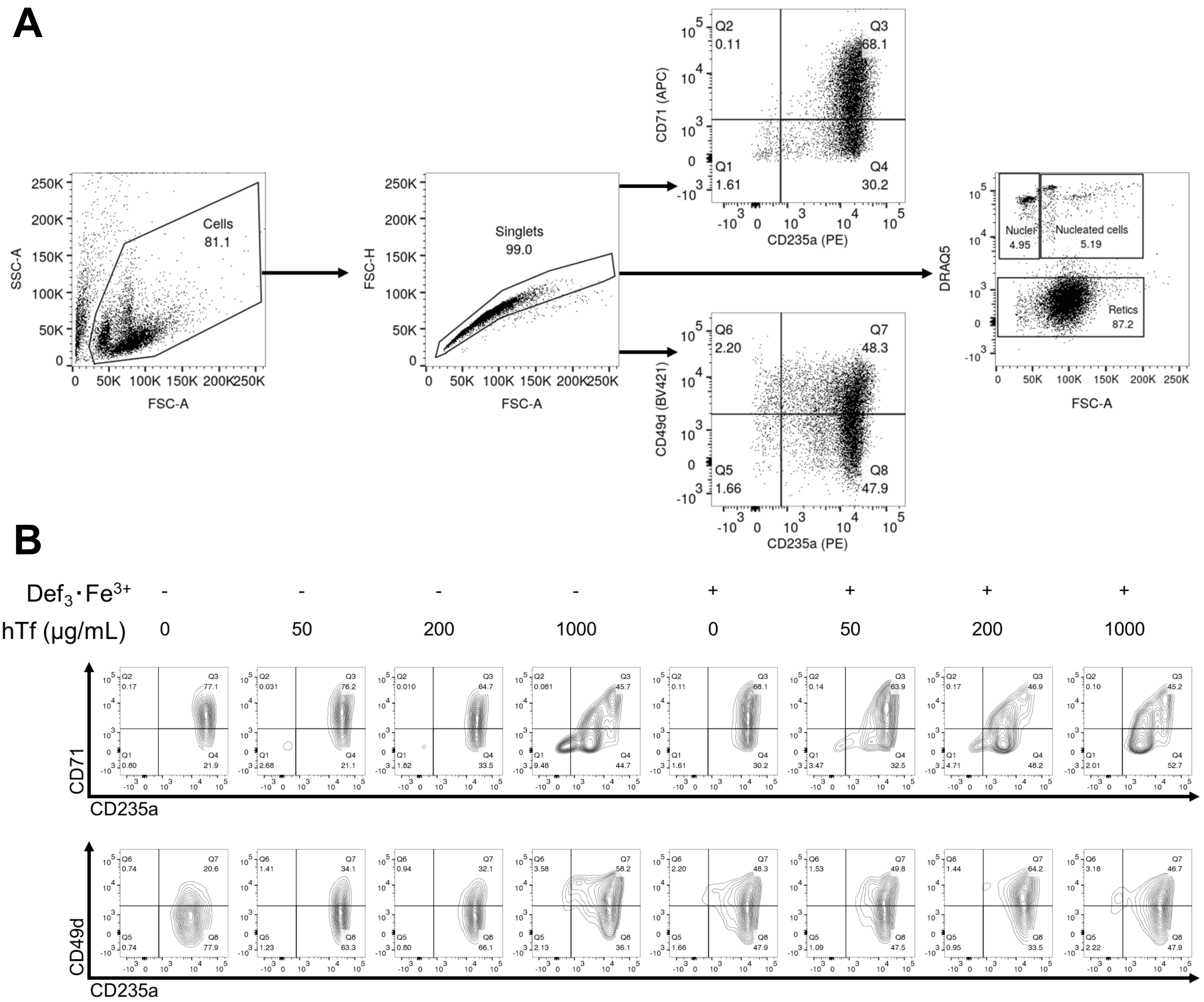

### Supplemental Figure 4

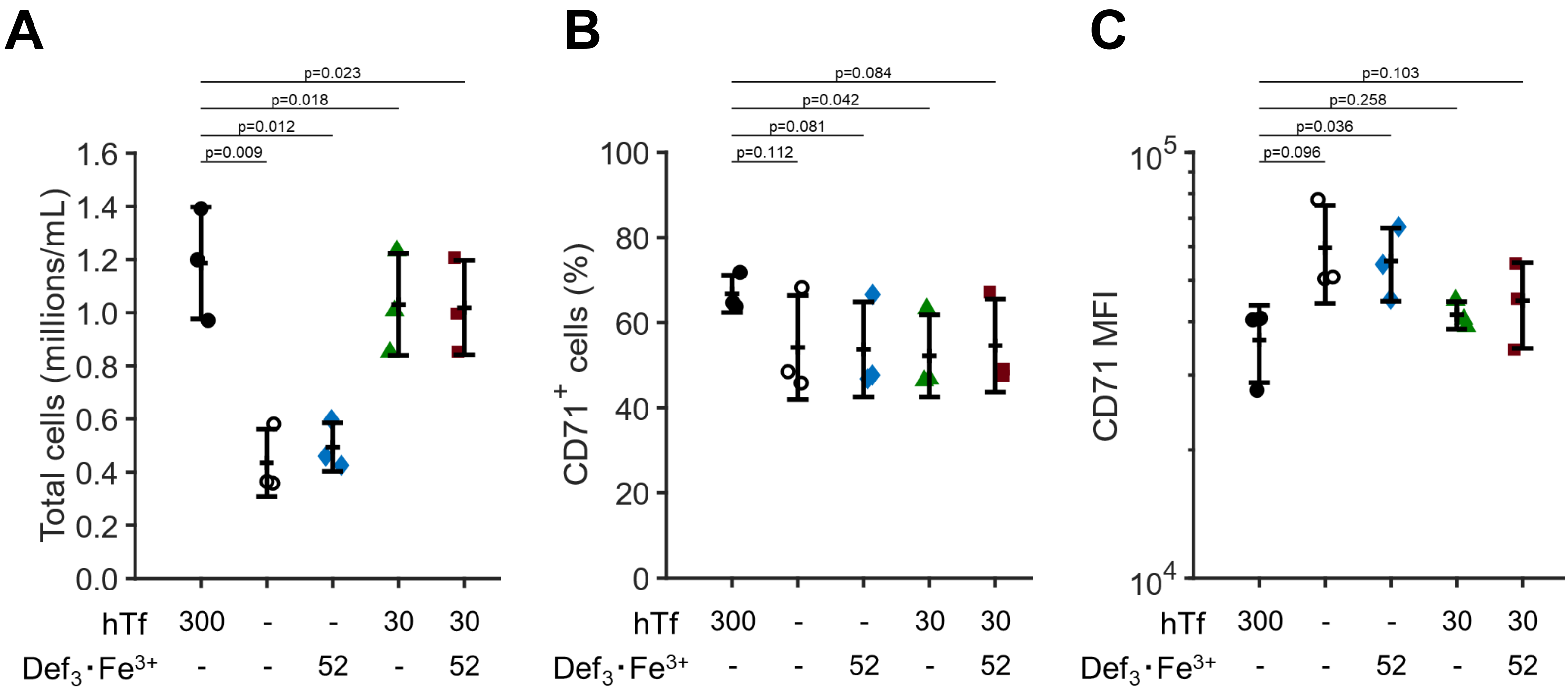

### Supplemental Figure 5

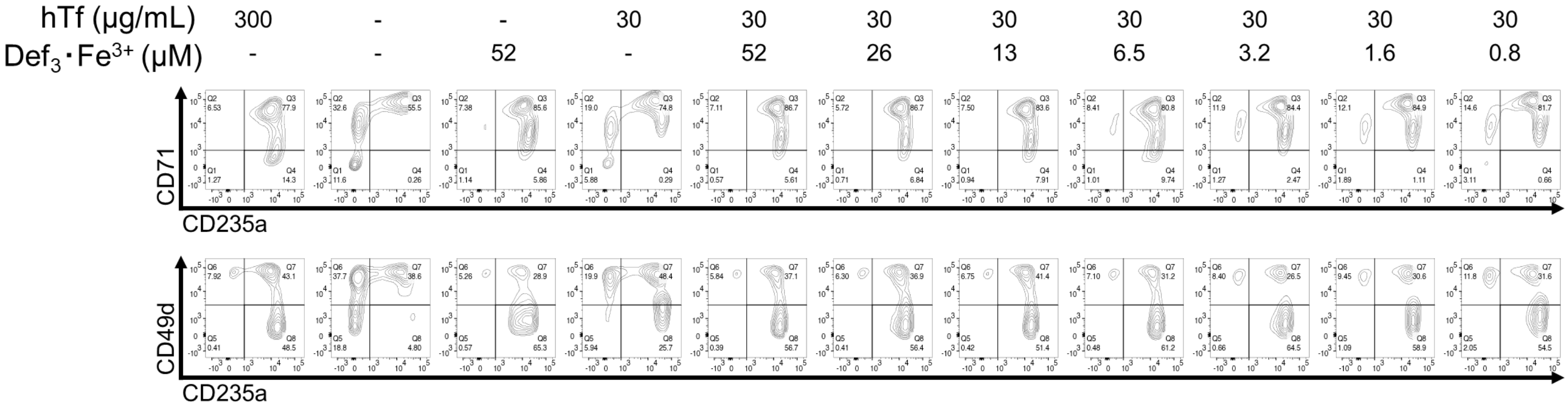

### Supplemental Figure 6

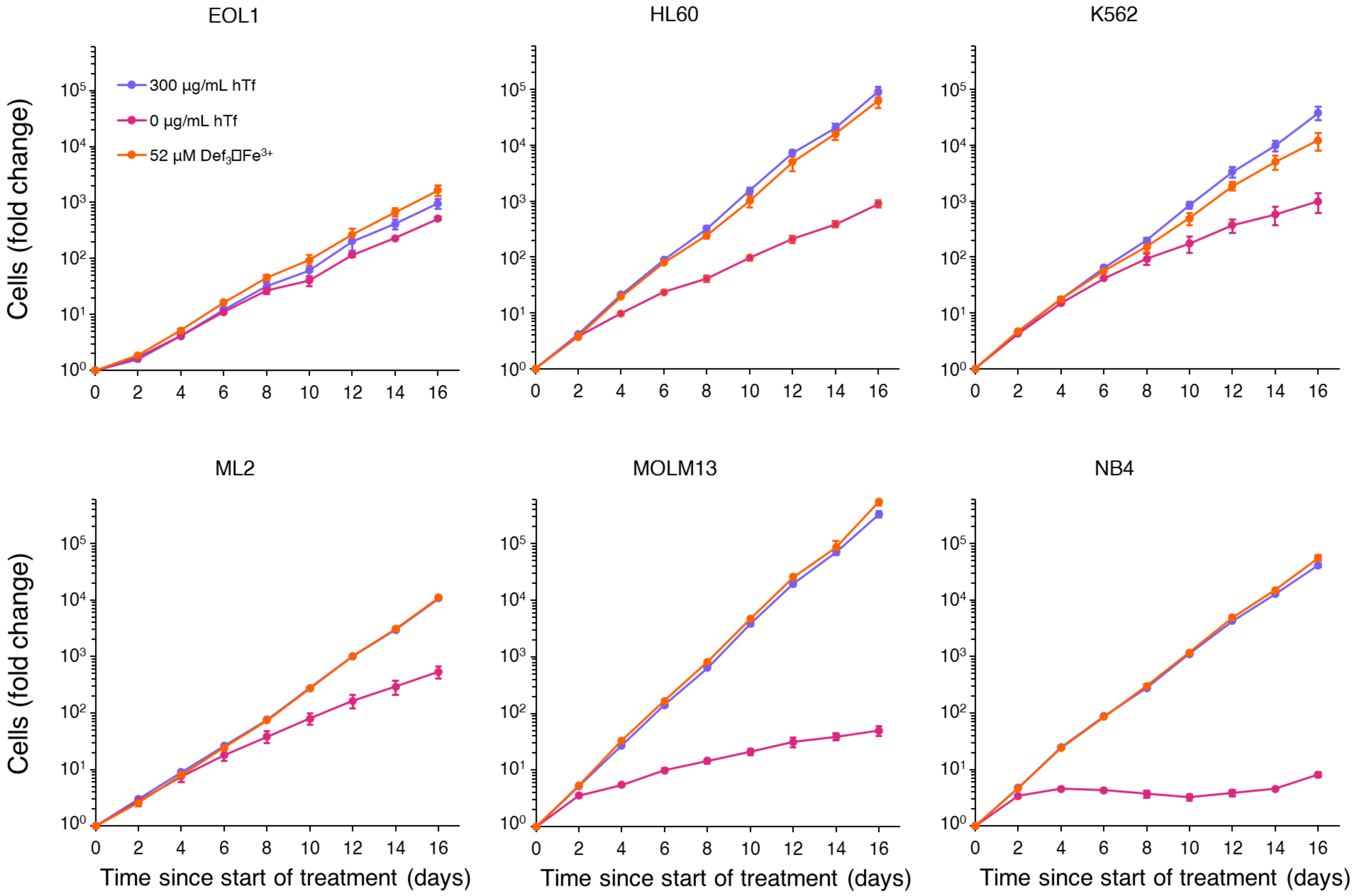
